# Insight into glycosphingolipid crypticity: Crystal structure of the anti-tumor antibody 14F7 and recognition of NeuGc GM3 ganglioside

**DOI:** 10.1101/2020.09.18.294777

**Authors:** Kaare Bjerregaard-Andersen, Hedda Johannesen, Fana Abraha, Aleksandra Šakanović, Daniel Groβer, Ünal Coskun, Gregor Anderluh, Stefan Oscarson, Ernesto Moreno, Michal Grzybek, Ute Krengel

**Affiliations:** Department of Chemistry, University of Oslo, NO-0315 Oslo, Norway; Department of Biosciences, University of Oslo, NO-0316 Oslo, Norway; School of Chemistry, University College Dublin, Belfield, Dublin 4, Ireland; Department of Molecular Biology and Nanobiotechnology, The National Institute of Chemistry, 1000, Ljubljana, Slovenia; Paul Langerhans Institute Dresden of the Helmholtz Zentrum Munich at the University Hospital and Faculty of Medicine Carl Gustav Carus of Technische Universität Dresden, Dresden, Germany; German Center for Diabetes Research (DZD e.V.), Neuherberg, Germany; Facultad de Ciencias Básicas, Universidad de Medellín, Medellín, Colombia; Kaare Bjerregaard-Andersen, H. Lundbeck A/S, Valby, Denmark; Fana Abraha, Recipharm OT Chemistry, Uppsala, Sweden

**Keywords:** anti-tumor antibody, carbohydrate-antibody interactions, carbohydrate-carbohydrate stacking, gangliosides, glycosphingolipid crypticity, lipid rafts, liposomes, *N*-glycolyl GM3, protein-carbohydrate interactions, X-ray crystal structure

## Abstract

Tumor-associated glycolipids such as NeuGc GM3 are auspicious molecular targets in antineoplastic therapies and vaccine strategies. 14F7 is an anti-tumor antibody with high clinical potential, which has extraordinary specificity for NeuGc GM3, but does not recognize the very similar, ubiquitous NeuAc GM3. Here we present the 2.3 Å crystal structure of the 14F7 binding domain (14F7 scFv) in complex with the NeuGc GM3 trisaccharide. Intriguingly, a water molecule appears to shape the specificity of 14F7. Using model membrane systems, we show that 14F7 recognizes NeuGc GM3 only above lipid concentrations that are likely to form glycolipid-rich domains. This “all-or-nothing” effect was exacerbated in giant unilamellar vesicles and multilamellar vesicles, whereas no binding was observed to 100 nm liposomes, emphasizing that the 14F7–NeuGc GM3 interaction is additionally modulated by membrane curvature. Unexpectedly, adding NeuAc GM3 strongly increased binding affinity to NeuGc GM3-containing liposomes. This effect may be important for tumor recognition, where the ubiquitous NeuAc GM3 may enhance 14F7 binding to NeuGc GM3-expressing cancer cells.

## Introduction

Cancer cells differ from healthy cells by aberrant glycosylation patterns, displaying tumor-associated carbohydrate antigens (TACAs)^1-3^. Immunotherapy offers the possibility of specifically targeting TACAs with high affinity through structure-based engineering of monoclonal antibodies^4-6^. The monoclonal antibody (mAb) 14F7 is an IgG_1_ raised by immunizing a BALB/c mouse with *N*-glycolyl GM3 (NeuGc GM3) complexed with very low-density lipoproteins (VLDLs)^7^. This antibody is known for its exquisite specificity and high affinity to NeuGc GM3, determined by ELISA to be in the low nanomolar range^7-9^. 14F7 has been used to verify the presence of the NeuGc GM3 in a range of tumors including retinoblastoma^10^, non-small cell lung cancer^11^, colon cancer^12^, breast cancer^7,13^ and melanoma^7^. Humanizing the mAb yielded 14F7hT (here referred to as 14F7 mAb), which retained its original ability to induce antibody-dependent cellular cytotoxicity in both human and murine NeuGc GM3-expressing cells^14,15^. 14F7 mAb has been reported to kill primary tumor cells by a complement-independent mechanism^16,17^, however, the details of its mode of action are unknown.

The ability of 14F7 to effectively differentiate between the highly similar NeuGc and NeuAc epitopes is intriguing. In fact, the two glycolipids only differ by the presence of one additional oxygen atom (H to OH) present in NeuGc GM3 (Figure 1A). Mutational studies have highlighted key residues involved in NeuGc binding^18^. The structural basis of the discrimination between NeuGc and NeuAc GM3 has, however, remained elusive. Partial understanding has been gained through the crystal structure of the 14F7 Fab^19^ and the more recent structure of a 14F7-derived single-chain variable fragment (scFv) harboring an alternative light chain^9^. Both 14F7 formats feature a long CDR H3 loop, which exhibits key residues for antigen binding.

**Figure 1.**
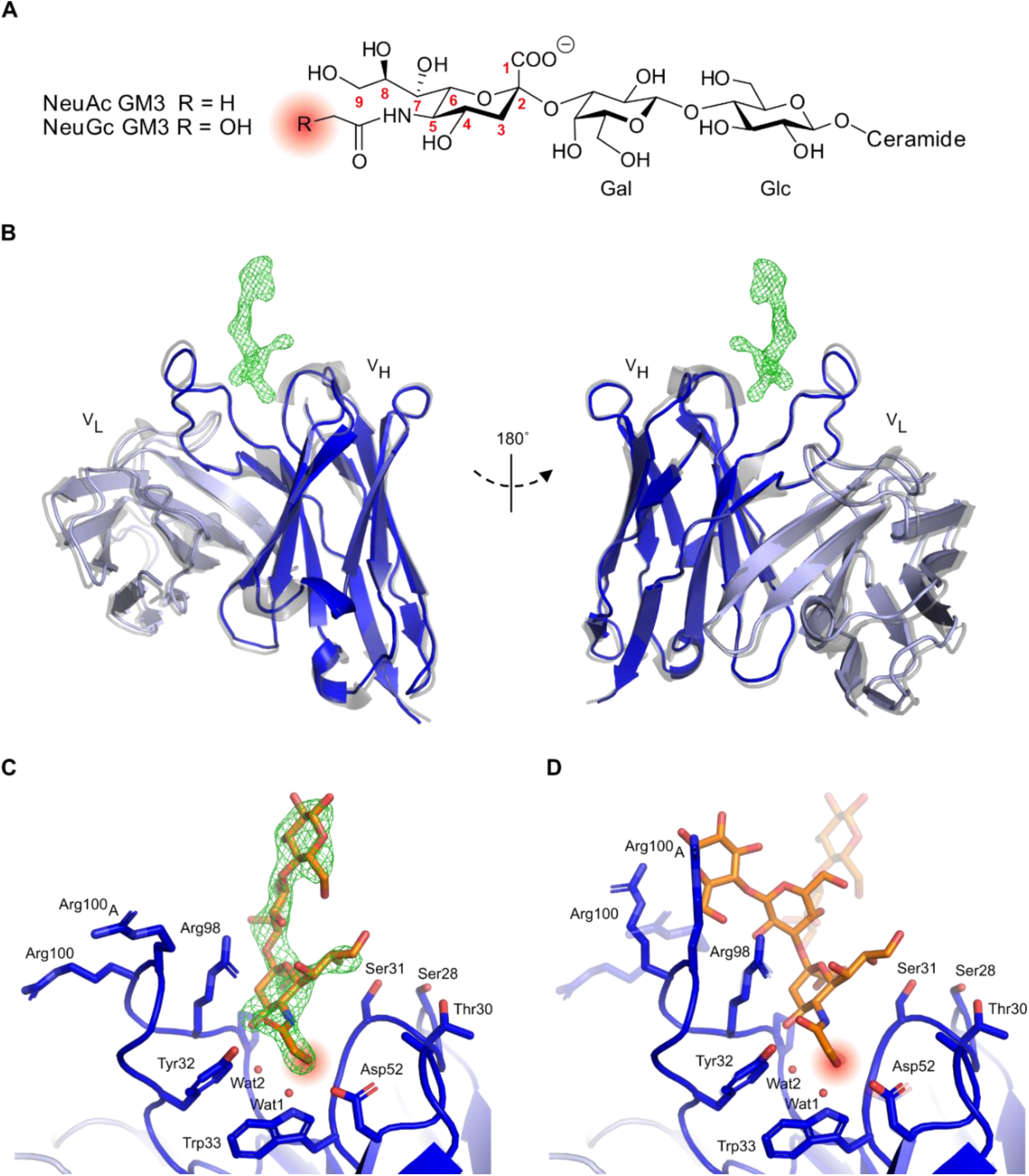
14F7 scFv complex with NeuGc GM3 trisaccharide. **A** Ganglioside structure. Neu5Gc GM3 is a xeno-antigen with a very similar structure to the common cellular glycolipid Neu5Ac GM3. The only difference consists of an additional oxygen atom in the *N*-glycolyl group of NeuGc compared to the *N*-acetyl group of NeuAc (highlighted in salmon) in the context of the trisaccharide Neuα2-3Galβ1-4Glcβ. **B** 14F7 scFv light (light blue) and heavy (dark blue) chains (PDB ID: 6S2I, chain A; this work). The 14F7 scFv apo-structure (PDB ID: 6FFJ^9^, chain A) is superimposed in grey. Difference electron density (*m****F*o**-*D****F*c**) for the carbohydrate ligand is shown at 3.0 σ (green mesh). **C** Structural model of 14F7scFv―NeuGc trisaccharide complex (synclinal conformation). Key amino acid residues and water molecules interacting with the glycan (orange) are labeled. **D** Alternative conformation of NeuGc GM3 trisaccharide with anticlinal glycosidic linkage between NeuGc and Gal (modeled), which buries a larger surface area on 14F7 scFv compared to the synclinal conformation observed in the crystal structure (transparent). Panel A was prepared with ChemDraw, panels B-D with PyMOL 2.2.0.

NeuGc GM3 is composed of a ceramide tail, buried in the plasma membrane of the cell, and an exposed trisaccharide head group featuring the sialic acid NeuGc at its tip^20^. While NeuGc GM3 is expressed in most mammals, it is absent from healthy adult human cells due to a partial deletion in the cytidine monophosphate-*N*-acetylneuraminic acid hydroxylase (*CMAH*) gene converting NeuAc to NeuGc^21,22^. However, dietary uptake of NeuGc GM3, *e*.*g*., from meat, can lead to low levels present in healthy tissue^23-27^. This in turn leads to a low level of autoantibodies, and NeuGc GM3 is therefore referred to as a xeno-autoantigen^28^. In contrast to NeuGc GM3, its *N*-acetyl counterpart is found ubiquitously in human cells and plays a role in control of numerous cellular signaling pathways^29,30^. NeuAc GM3 has also been shown to interact with integral membrane proteins, such as the insulin receptor^31^, or the epidermal growth factor receptor^32,33^. By mechanisms that are not yet well understood^23,28,34-37^, NeuGc GM3 is displayed to a larger extent by certain cancer cells and thus represents an attractive TACA.

Here we present the X-ray crystal structure of the scFv–NeuGc complex, elucidating the molecular basis for its discrimination between NeuAc and NeuGc GM3. Analysis of the crystal structure has been expanded through molecular modeling to propose an alternative binding mode of the GM3 lactose moiety that better explains previous mutagenesis data. Furthermore, our binding experiments of the 14F7 mAb and scFv to NeuGc GM3 reconstituted in liposomes show that the antibodies efficiently recognize the ganglioside only at high concentrations. Interestingly, the presence of NeuAc GM3 potentiates antibody recognition of NeuGc GM3, suggesting that 14F7 mAb and scFv can be potent tools for targeting low molar concentrations of the NeuGc GM3 antigen in NeuAc GM3-expressing cells.

## Results

### Crystal structure of 14F7 scFv in complex with NeuGc GM3 trisaccharide

The structure of the 14F7 scFv in complex with the NeuGc GM3 trisaccharide was determined to 2.3 Å resolution from a single trisaccharide-soaked crystal. Data collection and refinement statistics are summarized in Table 1. The crystal was obtained from the same batch of crystallization setups that earlier yielded the scFv apo-structure (PDB ID: 6FFJ^9^) and retained *P*2_1_ symmetry upon soaking, with similar unit cell parameters and four scFv molecules in the asymmetric unit. Two of the four molecules (chains A and B) were well defined by electron density in the CDR regions and could be modeled without breaks, whereas parts of CDR H3 could not be traced in chains C and D. One of the molecules (chain A) contained additional electron density corresponding to the trisaccharide ligand (Figure 1B). Inspection of the ligand complex revealed that only the sialic acid component (NeuGc) of the trisaccharide interacts with the antibody (Figure 1C), whereas the glucose moiety extends outwards towards the solvent, where it makes contacts with residues of a neighboring scFv within the same crystallographic asymmetric unit. In this binding mode, the glycosidic linkage between NeuGc and Gal adopts a synclinal conformation^38^.

**Table 1.**
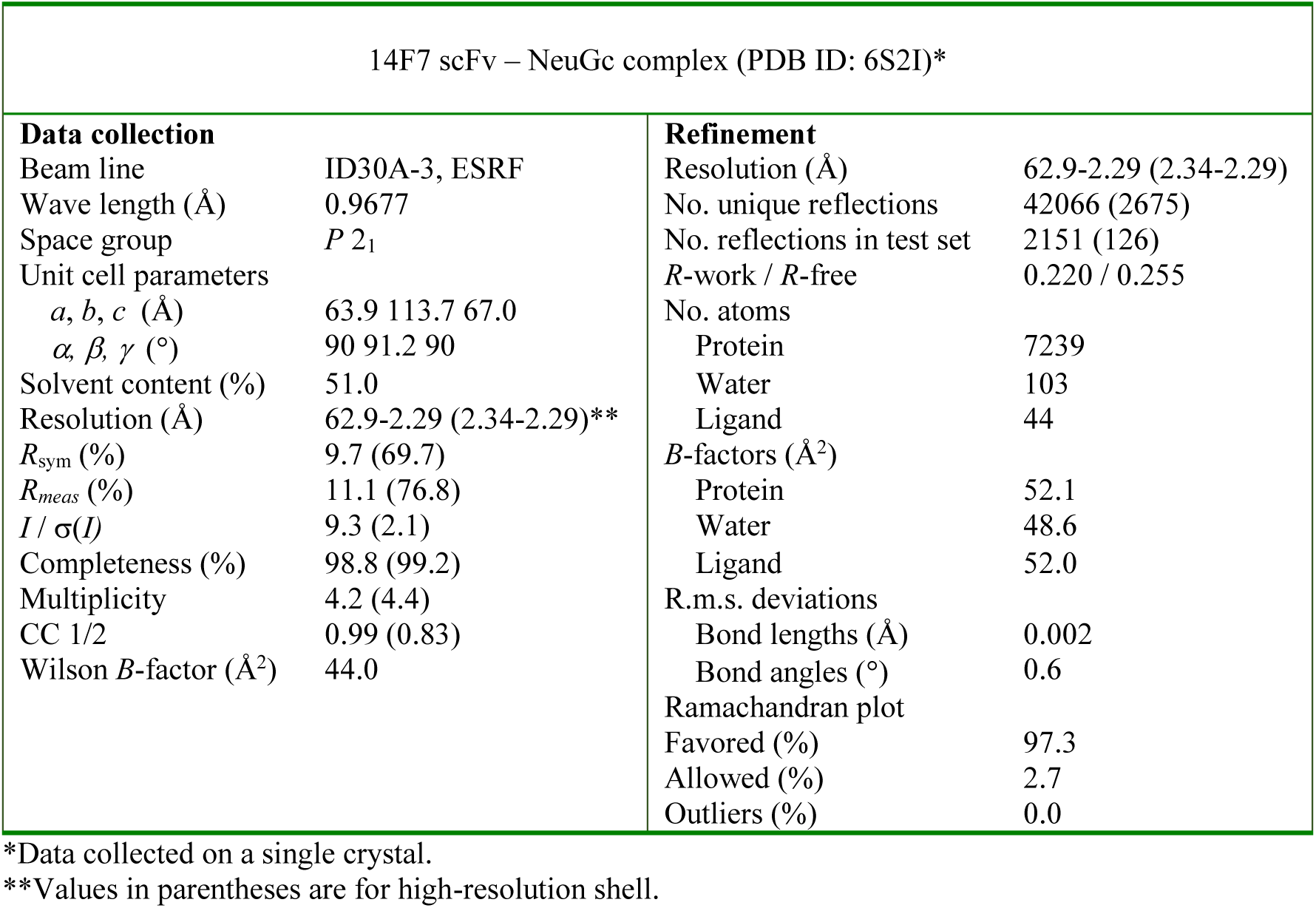
Crystallographic data collection and refinement statistics

| 14F7 scFv – NeuGc complex (PDB ID: 6S2I)* |  |  |  |
| --- | --- | --- | --- |
| Data collection |  | Refinement |  |
| Beam line | ID30A-3, ESRF | Resolution (Å) | 62.9-2.29 (2.34-2.29) |
| Wave length (Å) | 0.9677 | No. unique reflections | 42066 (2675) |
| Space group | *P* 2<sub>1</sub> | No. reflections in test set | 2151 (126) |
| Unit cell parameters |  | *R*-work / *R*-free | 0.220 / 0.255 |
| *a*, *b*, *c* (Å) | 63.9 113.7 67.0 | No. atoms |  |
| *α*, *β*, *γ* (°) | 90 91.2 90 | Protein | 7239 |
| Solvent content (%) | 51.0 | Water | 103 |
| Resolution (Å) | 62.9-2.29 (2.34-2.29)** | Ligand | 44 |
| *R*<sub>sym</sub> (%) | 9.7 (69.7) | *B*-factors (Å<sup>2</sup>) |  |
| *R*<sub>meas</sub> (%) | 11.1 (76.8) | Protein | 52.1 |
| *I* / *σ*(*I*) | 9.3 (2.1) | Water | 48.6 |
| Completeness (%) | 98.8 (99.2) | Ligand | 52.0 |
| Multiplicity | 4.2 (4.4) | R.m.s. deviations |  |
| CC 1/2 | 0.99 (0.83) | Bond lengths (Å) | 0.002 |
| Wilson *B*-factor (Å<sup>2</sup>) | 44.0 | Bond angles (°) | 0.6 |
|  |  | Ramachandran plot |  |
|  |  | Favored (%) | 97.3 |
|  |  | Allowed (%) | 2.7 |
|  |  | Outliers (%) | 0.0 |

\*Data collected on a single crystal.
\*\*Values in parentheses are for high-resolution shell.

Overall, the structure of the scFv–NeuGc GM3 trisaccharide complex is highly similar to the previously published scFv apo-structure^9^, with an average r.m.s.d. value of 0.6 Å for Cα atoms indicating very little structural change upon binding (Figure 1B). Also, the side chain conformations of amino acid residues in proximity of the saccharide binding site are very similar between the scFv complex and the apo-structure. Tyr32 and Tyr100_D_, both in direct contact with the ligand through H-bonds, shift by approximately 1 Å to accommodate binding. Most noticeably, Arg98 adopts a new conformation upon ligand binding, where it stacks against the sialic acid residue of the NeuGc GM3 trisaccharide (Figure 1C and D).

### Structural basis for 14F7 discrimination between NeuGc and NeuAc GM3

The interactions between 14F7 and the NeuGc GM3 trisaccharide are shown in Figure 2A and listed in Table 2. 14F7 has repeatedly been shown to strongly differentiate between NeuGc and NeuAc GM3 *in vitro, e*.*g*., probed by ELISA^7,9^. Therefore, the key determinant for discrimination must be found in the trisaccharide head group, where the only difference is the presence of an additional hydroxyl group in the *N*-glycolyl moiety of the sialic acid. Intriguingly, the *N*-glycolyl hydroxyl group does not itself provide any direct interaction with the scFv, except for a backbone interaction with Tyr32, but manifests its presence through a water molecule (Wat1; Figure 2A). Wat1 is part of a hydrated pocket coordinated by Trp33 and is also present in the 14F7 scFv apo-structure (PDB ID: 6FFJ^9^), thus it may be regarded as an extension of CDR H1. Wat1 not only interacts with the *N*-glycolyl hydroxyl group of NeuGc, but also with its 4-OH group, *via* a second water molecule (Wat2), which binds to the backbone oxygen of Ser96. On the protein side, Wat1 establishes an H-bond with the backbone NH of Trp33 and a weaker, out-of-plane H-bond with the aromatic π face of its indole pyrrole ring. Mutagenesis of Trp33 reveals that specificity is only maintained when this residue is exchanged by another aromatic residue, *i*.*e*., Phe and Tyr^18^. Especially the possible replacement by Phe emphasizes the importance of the aromatic interaction with Wat1. This trisaccharide-water complex, unable to form with NeuAc, places itself like a cassette into the bottom of the binding pocket formed by the backbone and side chains of Ser31, Tyr32, Pro97, Arg98 and Tyr100_D_. The difference in energetic contribution to binding of this exciting water-mediated ligand binding site remains to be explored.

**Table 2.**
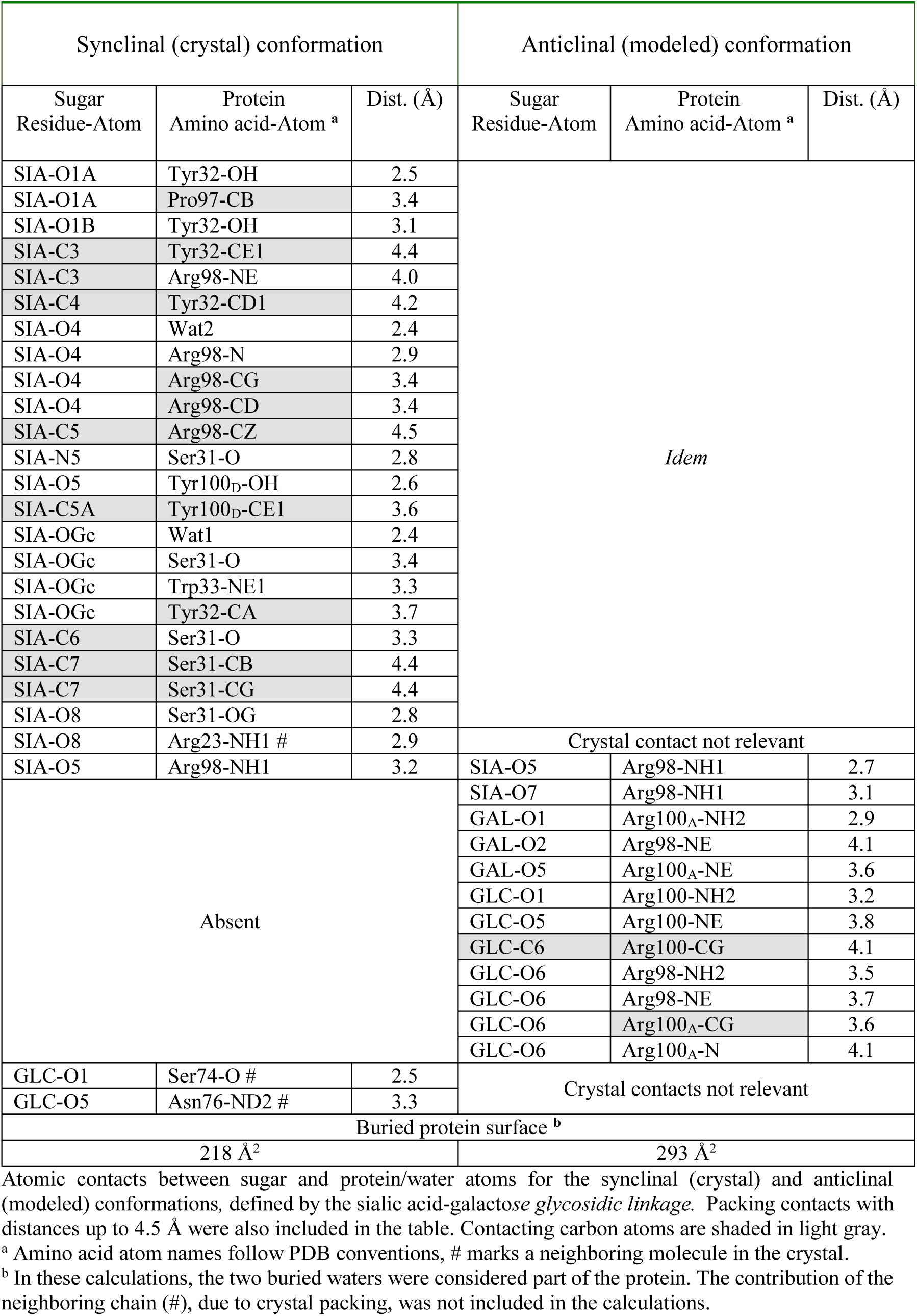
Protein-carbohydrate interactions.

**Figure 2.**
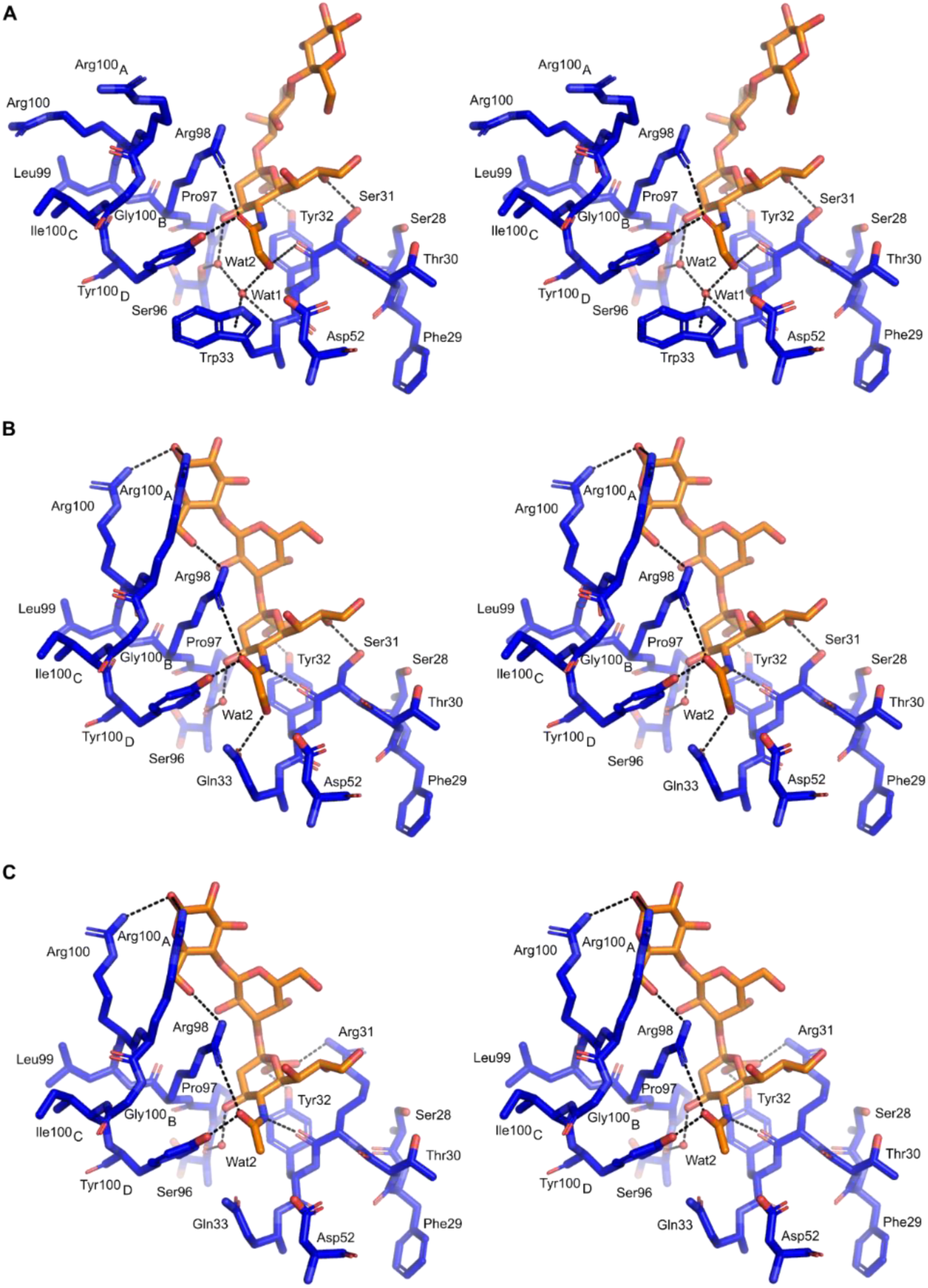
Stereo pictures showing the specificity of Trp33, W33Q and W33Q/S31R 14F7 variants in complex with NeuGc or NeuAc. **A** Crystal structure of 14F7 Trp33 (blue) bound to NeuGc (orange) in its experimentally determined conformation (PDB ID: 6S2I, chain A; this work). **B** Model of 14F7 W33Q variant, with NeuGc in the *in silico*-optimized anticlinal conformation. **C** Model of the cross-reactive 14F7 S31R/W33Q variant, with NeuAc in the anticlinal conformation. The figure was prepared with PyMOL 2.0.0.

### Alternative trisaccharide binding mode

In the crystal, NeuGc GM3 adopts synclinal torsion angles between NeuGc and Gal. In solution, a common alternative conformation of the NeuGc GM3 trisaccharide has an anticlinal glycosidic linkage. Reasoning that crystal packing might have forced the orientation of the lactose moiety of NeuGc GM3 into the conformation observed in the crystal structure, we modeled an alternative binding mode for the trisaccharide, where NeuGc remained exactly as in the crystal structure, but the two torsion angles of its glycosidic linkage with galactose adopt the anticlinal conformation (Figure 1D). We found that this binding geometry, which is hindered by the crystal packing, brings additional favorable contacts between the trisaccharide and CDR-H3, including interactions between both Arg100 and Arg100_A_ with the trisaccharide glucose residue (Table 2). It also increases the buried surface area by more than one third, from 218 Å^2^ to 293 Å^2^. Furthermore, in this binding mode, Arg98 becomes more tightly packed against the trisaccharide (Figure 1D, Figure 2AB). This is in good agreement with mutagenesis data showing the critical role of this amino acid, which did not tolerate any substitution^18^.

### Models of 14F7 variants explore functional mapping data

In previous work, we used phage display to perform extensive mutagenesis studies on the 14F7 heavy chain CDRs^18^. These studies identified several positions in CDRs H1 and H3 as important for recognizing NeuGc GM3, *e*.*g*. Trp33, Asp52, Arg98, Arg100, Arg100_A_ and Tyr100_D_. In addition we found that several single residue substitutions, yielding *e*.*g*., S28R, T30R, S31R and W33Q conferred different levels of cross-reactivity to the antibody; and some double or triple combinations even raised the affinity to NeuAc GM3 to the same level as for NeuGc GM3^18^. Here we modeled one of these variants (W33Q) in complex with NeuGc GM3 (Figure 2B) and another (S31R/W33Q) in complex with NeuAc GM3 (Figure 2C), in order to interpret the mutagenesis data. The introduction of an arginine residue in the antigen binding site is likely to yield a salt bridge with the sialic acid carboxylate. Gln33 (as in W33Q) probably interacts directly with the *N*-glycolyl OH of NeuGc GM3, replacing Wat1 (Figure 2B).

### 14F7 binding to NeuGc GM3 occurs only at high glycolipid densities

To better understand the mode of interaction between 14F7 and NeuGc GM3 in the lipid bilayer, we performed a series of binding experiments using different model membrane systems. First, we chemically labeled 14F7 mAb and scFv with fluorescent dyes and tested their binding to giant unilamellar vesicles (GUVs) containing various amounts of NeuGc or NeuAc GM3 in a background of DOPC and cholesterol (Figure 3). Below 2 % NeuGc GM3, no binding to GUVs was detected. Even upon increasing the glycolipid concentration from 2 % to 5 % NeuGc GM3, only ∼3 % of all vesicles showed antibody binding. However, when 10 % NeuGc GM3 was used for GUV formation, all vesicles were labeled (Figure 3), suggesting that 14F7 mAb is not capable of recognizing individual NeuGc GM3 glycolipids as antigens. Upon surpassing a critical glycolipid density threshold, however, antigen recognition becomes highly efficient. In contrast, no binding was observed for 10 % NeuAc, confirming the specificity of 14F7 (Figure 3).

**Figure 3.**
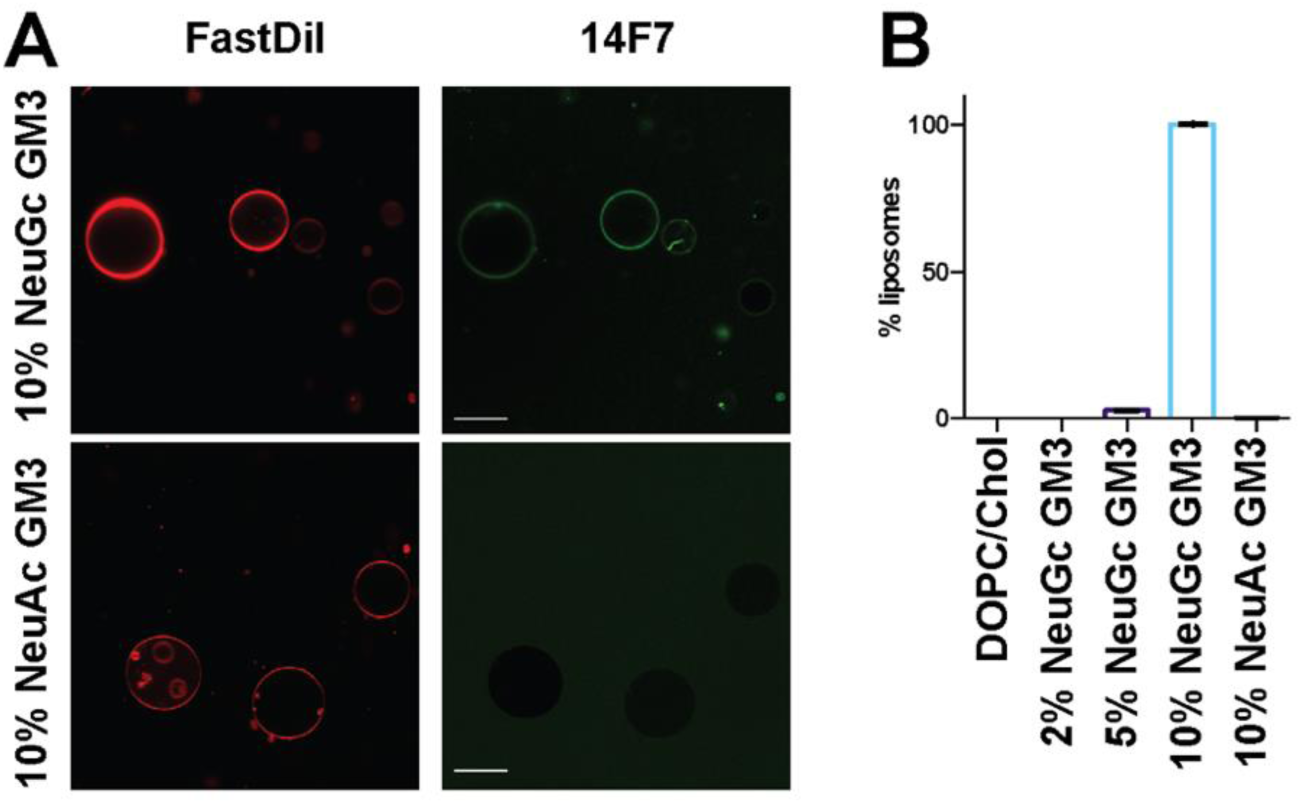
Binding of 14F7 to GUVs composed of DOPC/Chol/GM3. **a**14F7 was chemically labeled with Dylight-488 (green) and was added to GUVs containing FastDiI as a membrane marker (red). The scale bar corresponds to 20 μm. **b** Percentage of GUVs showing binding of 14F7-488. Representative images for all lipid compositions are shown in Figure S1.

The experiment was repeated with fluorescently labeled 14F7 scFv instead of 14F7 mAb, however, no binding was observed, even at high NeuGc concentrations. To test whether this failure in binding was related to loss of function in the 14F7 scFv or an artifact of chemical labeling, we performed differential-scanning calorimetry (nanoDSF) of the unlabeled and labeled scFv fragment (Figure S3). After chemical labeling with Dylight488-NHS, the T_m_ of 67 °C for the native 14F7 scFv flattened and shifted its maximum to 72 °C (Figure S3). In addition, the scattering of the solution increased substantially, suggesting that the sample becomes polydisperse. Both of these results indicate an overall change in the structure of the scFv upon labeling that probably also perturbs the binding site. We therefore opted for liposomal flotation experiments that do not require labeling of the protein.

### Membrane curvature can affect NeuGc GM3 presentation

To test if both 14F7 mAb and scFv can recognize NeuGc GM3-containing liposomes, we performed flotation assays with unlabeled protein. Large unilamellar vesicles (LUVs) have an advantage over GUVs in that they form more homogenous liposome populations due to their preparation method, using freeze-thaw cycles, followed by multiple extrusions through polycarbonate filters. After the flotation assay, the gradients were fractionated from top to bottom, and binding of 14F7 to vesicles was detected by Western blotting. Antibody binding to LUVs is indicated by its presence in the top, low-density fractions (fractions 2-4), otherwise, the protein would be pelleted at the bottom (fractions 10-11) (Figure 4A). In the initial experiment, LUVs containing 10 % NeuGc GM3 were used, but only a small portion of either 14F7 mAb or scFv was detected in the liposome-containing fractions, suggesting a very weak interaction with the LUVs (Figure 4BC). Thin layer chromatography (TLC) analysis of the liposomes recovered after flotation confirmed the presence of NeuGc GM3 in the vesicles, thus the weak binding was puzzling. The major difference between the LUVs and the liposomes used in the fluorescent study (GUVs) was their size (Figure 4D). Therefore, we repeated the flotation experiment using non-extruded, multilamellar liposomes (MLVs), which correspond in size to the GUVs. Interestingly, strong binding for both 14F7 mAb and scFv was observed for the MLVs, suggesting that membrane curvature plays an important role for antigen recognition (Figure 4BC). A comparative analysis of lipid composition of both LUVs and MLVs confirmed that in both cases the composition of the vesicles was the same (Figure 4E). In fact, the amount of NeuGc GM3 that was available for binding should be much higher in LUVs than in MLVs, as only the outermost membrane leaflet can be probed by the antibodies. When NeuAc GM3 was used in either LUVs or MLVs, no binding was observed, neither for 14F7 mAb nor for scFv, again confirming the high specificity of 14F7 (Figure S2).

**Figure 4.**
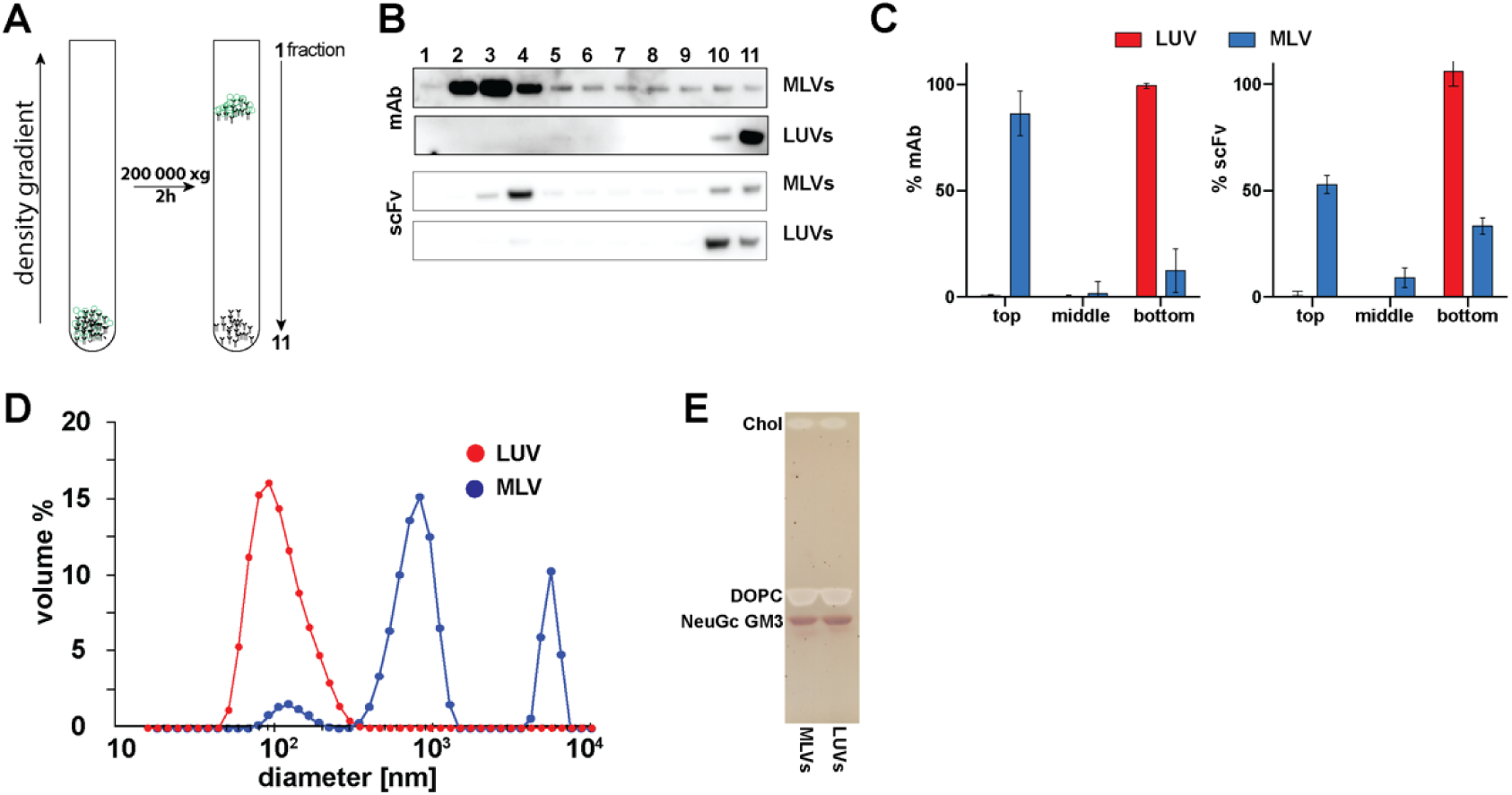
Binding of 14F7 to LUVs and MLVs composed of DOPC/Chol/NeuGc GM3 in flotation assays. **A** Setup of the LUV flotation assay. After the centrifugation, 11 fractions were collected from the top. Proteins bound to vesicles accumulate in fractions 2-4, while unbound proteins are in the bottom fractions (10-11) **B** Representative Western blot of fractions collected after flotation. **C** Binding was quantified using the AIDA software as follows: top (fractions 1-4), middle (5-8) and bottom (9-11). Error bars represent standard deviations of three independent experiments. **D** Dynamic light-scattering measurement of the size of the vesicles used in the flotation assay. **E** Thin layer chromatography (TLC) of lipids extracted from the vesicles after flotation (combined fractions 2-4). The lipids were stained with orcinol.

### NeuAc GM3 potentiates 14F7 binding to NeuGc GM3

We decided to probe 14F7 mAb and scFv binding to MLVs containing various NeuGc GM3 concentrations (0-10 %) by electrochemiluminescence immunoassay (EIA), and also tested combinations of NeuGc and NeuAc GM3. The *K*_D_ values for vesicles containing 10 % NeuGc GM3 were estimated to be approximately 34 nM and 3.4 *μ*M for 14F7 mAb and scFv, respectively by EIA (Figure 5AB), representing a large gain in apparent affinity (approximately 100-fold) for the mAb *versus* the scFv, in contrast to the previous ELISA studies^8,9^. Interestingly, NeuGc GM3 binding and recognition by 14F7 greatly increased when NeuAc GM3 was introduced into the same vesicles, *e*.*g*., binding of both 14F7 mAb and scFv to liposomes containing a mixture of 2 % NeuGc and 8 % NeuAc GM3 was significantly higher compared to liposomes containing only 2 % NeuGc. In fact, 14F7 scFv bound equally strongly to vesicles containing 5 % NeuGc or a 2/8 % mixture of NeuGc and NeuAc GM3. For 14F7 mAb, binding to the 2/8 % mixture even exceeded the binding efficiency observed for vesicles containing 5 % pure NeuGc GM3. This effect of potentiation was also observed for liposomes containing an equal molar ratio of NeuGc/NeuAc GM3 (5/5 %), which showed higher binding efficiency compared to vesicles that only contained 5 % NeuGc GM3 (Figure 5AB). Similar results were obtained by surface plasmon resonance (SPR) spectroscopy (Figure 5C). This demonstrates that not only the amount of NeuGc GM3, but also the overall GM3 ganglioside concentration affects the ability of 14F7 to recognize and bind NeuGc GM3.

**Figure 5.**
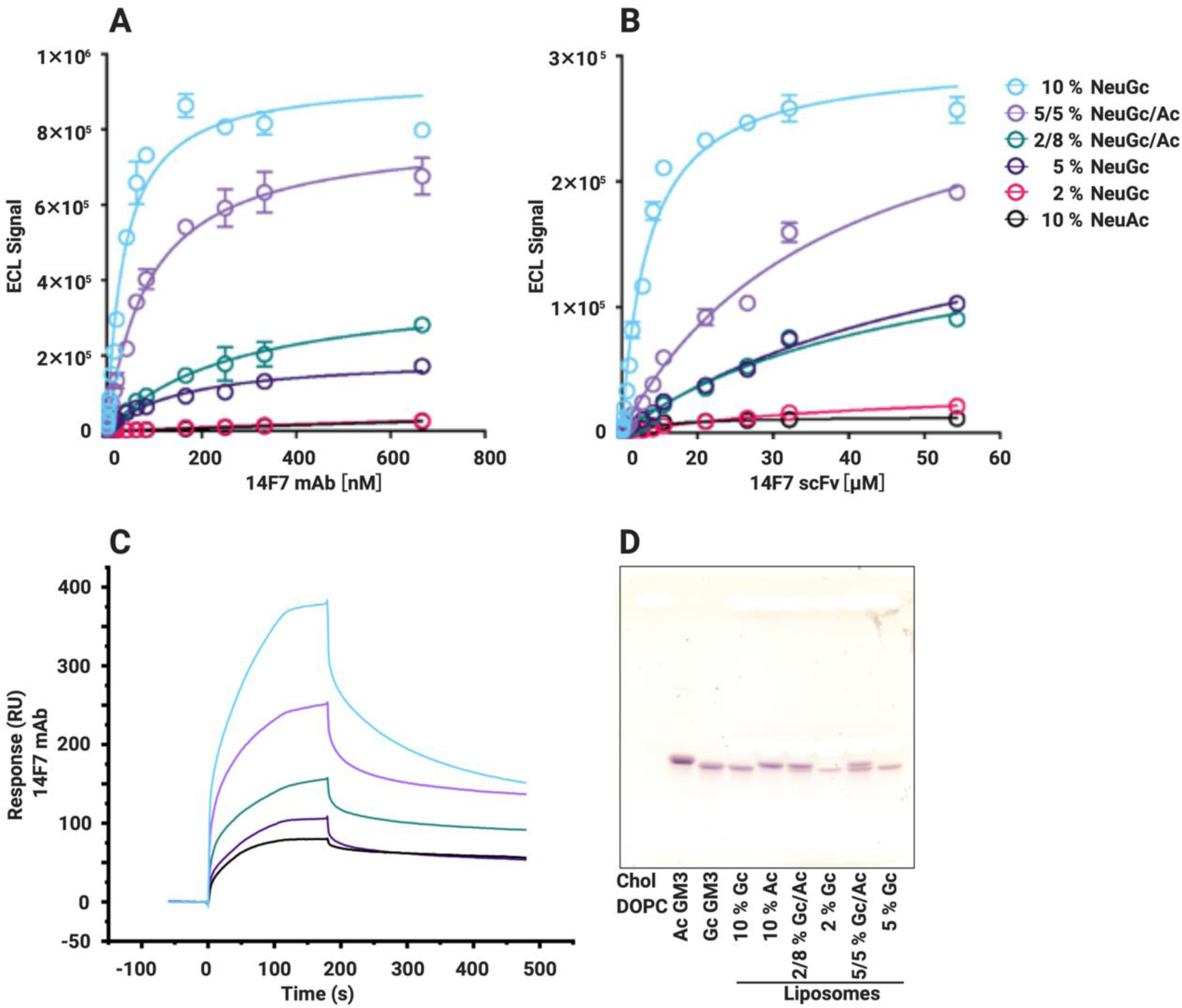
Addition of NeuAc GM3 increases binding efficiency of 14F7 to NeuGc GM3. Binding of (**A**) 14F7 mAb and (**B**) 14F7 scFv to MLVs composed of DOPC/Chol/GM3 measured by EIA. Representative plots of 3 independent measurements. **C** SPR spectroscopy of 14F7 mAb binding to an HPA chip with a monolayer of DOPC/Chol/GM3. The sensorgram represents one of two independent measurements (to save scarce sample), resulting in the same trend observed by EIA. **D** Thin layer chromatography of lipids extracted from the vesicles used in EIA assay. NeuAc and NeuGc GM3 were abbreviated Ac and Gc, respectively. The lipids were stained with orcinol.

### Discussion

Gangliosides are sialic-acid containing glycosphingolipids present in the plasma membranes of all vertebrates. Together with cholesterol, sphingomyelin and specific membrane proteins, they are concentrated in membrane nanodomains often referred to as lipid rafts^39-45^. Gangliosides are functionally important and are known to modulate cellular signaling^46-49^. Despite decades of studies, the structure and function of these cell surface antigens remain to be fully appreciated, and only few anti-ganglioside antibodies have been raised^50^.

### 14F7 specificity

14F7 can distinguish the very small chemical difference between the gangliosides NeuGc and NeuAc GM3^7,9^, and even more remarkably, we have now discovered that it does so indirectly, through a water molecule. NeuGc GM3 engages in two water-mediated interactions with Trp33, one with its main chain and one with the π-system of the indole side chain (both through Wat1; Figure 2A). Such an interaction is weaker than an ordinary hydrogen bond^51^, however, the importance of this interaction is highlighted by the fact that substitution of Trp33 with Phe or Tyr maintains specificity, while non-aromatic residues abolish binding or allow cross-reactivity with NeuAc GM3^18^. H-bonds commonly mediate specificity in antibody-antigen recognition through direct contact between paratope and epitope side chains^52^. In the case of 14F7, Wat1 is already present in the protein apo-structure (PDB ID: 6FFJ^9^). A thorough analysis of water–tryptophan interactions indicates that the six-membered ring of the indole side chain favors π-OH interaction, while the five-membered pyrrole ring favors π-lone pair interaction^53^. The latter appears to be the case for Wat1, thus positioning it as an H-bond donor for the *N*-glycolyl group of NeuGc GM3. While it is well known that the hydration shell is important for protein structure and function^54,55^, including the recognition of carbohydrates^56-58^ and antibody-antigen interactions^59-62^, the complexity of antibody engineering is highlighted by our finding of this indirect, water-mediated specificity.

### Selectivity *versus* cross-reactivity

NeuGc is bound to the bottom of a cleft formed by the variable heavy chain of 14F7 (Figure 2A), which is separated from the variable light chain through the long CDR H3 loop. The predicted NeuGc recognition site has previously been functionally mapped by a combinatorial phage display strategy using an alternative format of 14F7 scFv^18^. The study revealed that substitution of Trp33 in combination with residues 28, 30 or 31 could yield cross-reactive 14F7 variants (*e*.*g*., S28R/S30R/W33Q, S31R/W33Q and S28R/S31R, and to a lesser extent by single amino acid substitutions)^18^. Therefore cross-reactivity is likely mediated through direct interactions with the sialic acid residue, in particular by a salt-bridge to the negatively charged carboxylate group found in both NeuGc and NeuAc GM3. To further explore the mutagenesis data, we modeled the 14F7 S31R/W33Q variant in complex with NeuAc GM3 (Figure 2C). Substituting Trp33 as in 14F7 W33Q likely leads to the replacement of Wat1 by the glutamine side chain amide, which can interact directly with the *N*-glycolyl OH of NeuGc GM3 (Figure 2B). This mutation alone decreased NeuGc GM3 binding, but promoted a weak interaction to the NeuAc variant of GM3^18^. Substitution of Ser31 with Arg (S31R) probably trades an H-bond to one of the NeuGc glycerol hydroxyls for a charge interaction of the guanidinium moiety with the sialic acid carboxyl group found in both NeuGc and NeuAc GM3 (Figure 2C), thus conferring some cross-reactivity to the antibody. Arginine substitutions of Ser28 (S28R) or Thr30 (T30R) likely elicit similar effects. Interestingly, in spite of this additional interaction, substituting Ser31 for Arg, either alone or combined with other amino acid substitutions, hardly increased the affinity for NeuGc GM3^18^.

Although it may seem counterintuitive that NeuAc could bind to a polar pocket, a polar environment is not unprecedented for NeuAc. For example, cross-reactive rotaviruses that recognize both NeuAc and NeuGc GM3 have been shown to display similar polar, water-containing pockets to accommodate the acetyl or glycolyl groups of their glycan receptors^63^. Favorable interactions elsewhere, *e*.*g*., with the sialic acid carboxylate or glycerol chain, may well compensate for less favorable interactions of the *N*-acetyl group. In fact, it is likely that selectivity of NeuGc over NeuAc GM3 requires a fine balance of interactions, and that too tight binding of the sialic acid residue may prevent selectivity and would tip the balance towards cross-reactivity towards NeuGc and NeuAc GM3.

### Glycan conformation and antibody recognition

In the crystal structure of the scFv–saccharide complex (PDB ID: 6S2I; this work), the saccharide adopts a synclinal conformation (Figure 1C), and the only interaction with 14F7 is *via* the sialic acid (Figure 2A). However, the carbohydrate conformation may be forced by the crystal, into which the ligand was soaked. For example, we note that the anticlinal conformation would lead to clashes with other protein molecules in the crystal, whereas the synclinal conformation is stabilized by an interaction of the sialic acid glycerol chain with V_L_ residue Arg23 of a neighboring scFv in the crystal (Table 2). In a biological context (and in solution), the saccharide would be free to adopt both conformations (Figure 1D) – also the anticlinal conformation, which provides a larger contact surface with the antibody (293 *versus* 218 Å^2^). Dynamic binding may in fact provide an entropic advantage. In both conformations, the glycosidic linkage between NeuGc and Gal places the key CDR H3 residue Arg98 in a central position for interaction with the NeuGc GM3 trisaccharide (Figure 2AB), explaining why any substitution of this residue renders it incompatible with binding. In anticlinal conformation, Arg98 can additionally interact with the glucose moiety of NeuGc GM3 through H-bonds. This is also true for Arg100 and Arg100_A_, which are located at the tip of CDR-H3. Moreover, the arginine residues exposed on CDR H3 create a strongly positively charged surface patch that will likely also interact with other components of the plasma membrane. The observation that these residues, in general, can be exchanged while maintaining a positive charge^18^, indicate non-specific interactions with the membrane through negative charges found in the proximity of the target antigen, such as other phospholipids, gangliosides or proteins.

### 14F7–membrane interactions: “All-or-nothing” effect

To better understand the mode of interaction between the 14F7 and NeuGc GM3 in the context of a membrane, we performed binding measurements using different membrane-mimetic systems. These studies confirmed the selectivity of 14F7. However, binding was only observed above a concentration threshold of the NeuGc GM3 (Figure 3B). The observed “all-or-nothing” effect for glycolipid recognition is not a new concept in itself. For example, Nores *et al*. observed that the murine mAb M2590 only recognized the GM3 antigen when the ganglioside concentration in a membrane reached a threshold of 8 %, as determined by binding to liposomes^64^. That 14F7 mAb recognizes NeuGc GM3 in a similar manner is intriguing and suggests that at low concentrations, the antigen remains “cryptic”. For mAbs, avidity immediately springs to mind as logical explanation for such a threshold effect, *i*.*e*., if recognition by both Fabs is required to detect and enhance binding. However, since the same effect is also observed for the 14F7 scFv, both related to density and curvature, this clearly indicates that avidity cannot be the main cause of the observed effect. Instead, it suggests that at low concentrations, the glycolipid conformation in the membrane does not allow recognition by the antibody (Figure 6A). At high concentrations, the gangliosides may pack differently, for example through carbohydrate stacking^65^, and expose their *N*-glycolyl group to enable recognition (Figure 6B). The surrounding sialic acid residues from other GM3 molecules close-by may further enhance affinity due to the increased negative charge. The fact that the addition of NeuAc GM3 to low concentrations of NeuGc GM3 enables and enhances 14F7 binding (Figure 5), supports this hypothesis.

**Figure 6.**
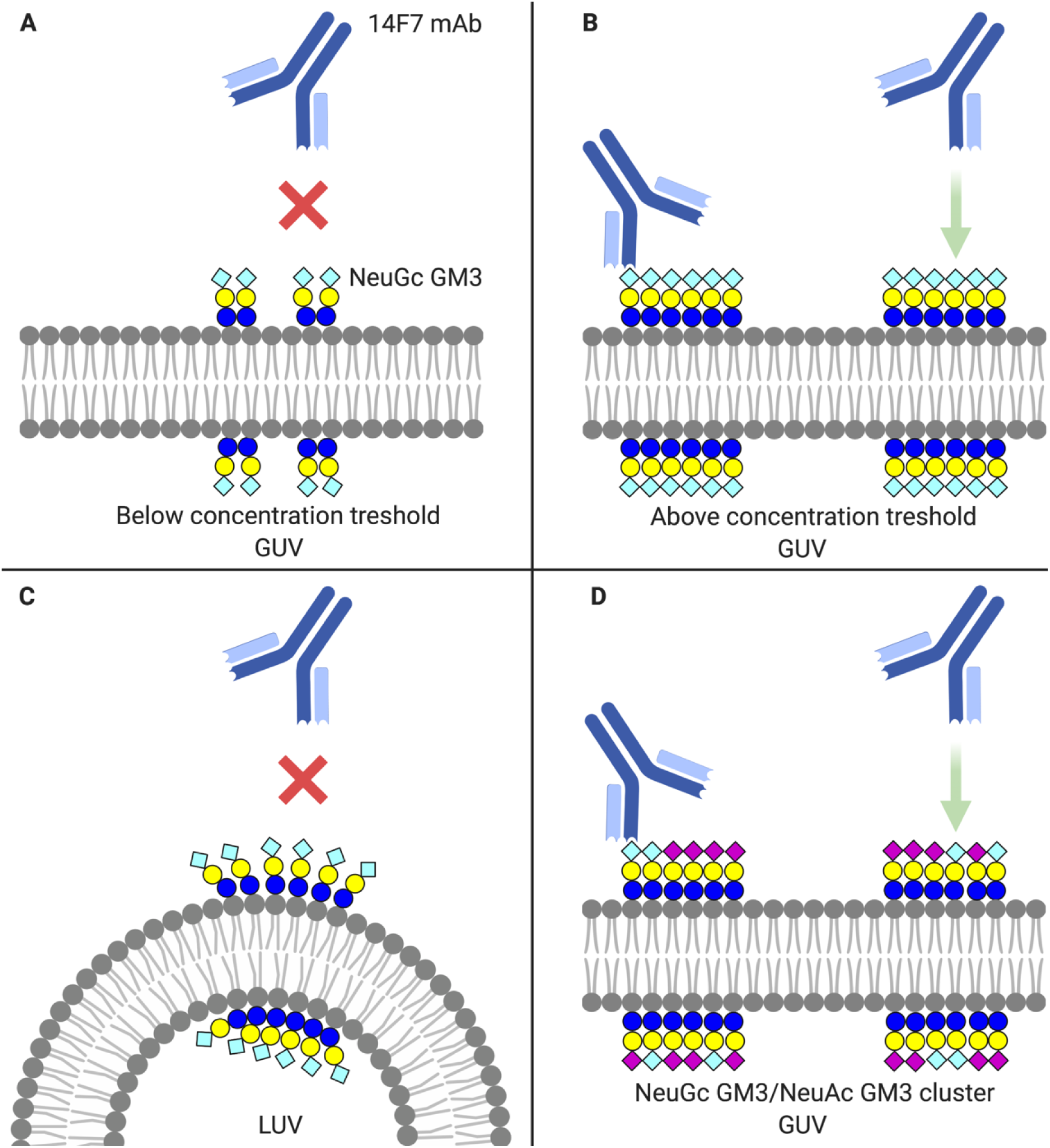
Model of NeuGc GM3 recognition by 14F7. **A** 14F7 binding is not observed at low NeuGc GM3 concentrations. **B** Above a ganglioside concentration threshold, 14F7 attaches to NeuGc GM3-containing glycolipid clusters on low-curvature membrane surfaces (GUV). **C** 14F7 does not bind to NeuGc GM3 in highly curved LUVs. **D** The addition of NeuAc GM3 to NeuGc GM3-containing liposomes enables 14F7 binding even at low NeuGc GM3 concentrations, possibly through the formation of functional glycolipid clusters. Carbohydrate symbols follow the nomenclature of the Consortium for Functional Glycomics: *N*-acetyl neuraminic acid – purple diamond; *N*-glycolyl neuraminic acid – light blue diamond; galactose – yellow circle; glucose – blue circle. Prepared with BioRender.

### NeuGc GM3 clustering

In contrast to the receptor-binding B-pentamer of the cholera toxin (CTB), which is commonly used to label GM1 molecules and can lead to ganglioside clustering^66^, 14F7 appears to bind only to pre-existing NeuGc GM3 assemblies and not drive the formation of such. This conclusion is based on the observation that (i) the monovalent scFv fragment showed similar binding characteristics as the mAb, and (ii) we never observed any domain formation in GUVs, even after overnight incubation with the divalent 14F7 mAb. For CTB, domain formation was observed when the protein was incubated with GUVs containing substantially lower amounts of its main glycolipid receptor, GM1^66^.

### Curvature effect

The collective behavior of lipids and physicochemical membrane properties can be directly modulated by temperature, pressure and molecular stress^67,68^. Another important parameter is membrane curvature^49,69-71^. Unexpectedly, we observed that the anti-tumor antibody 14F7 preferentially recognizes NeuGc GM3 present in GUVs or MLVs, but not in 100 nm LUVs. We hypothesize that high positive curvature might disrupt carbohydrate stacking interactions critical for exposure of the NeuGc *N*-glycolyl group, preventing NeuGc GM3 recognition as illustrated in Figure 6C.

Many lipids have a cone shape rather than a cylindrical shape^72^. In a flat lipid bilayer, the packing of cone-shaped lipids leads to packing tension (line-tension) that can be relieved by the spontaneous formation of higher-order submicron clusters^69,70^. This could be the driving force for NeuGc GM3 clustering, where ganglioside and cholesterol form liquid-ordered structures surrounded by liquid-disordered regions^73,74^. It would be interesting to test if also negatively curved interfaces (*e*.*g*., caveola and endocytic pits) elicit the preferable conformation for 14F7 recognition.

### New applications for 14F7?

In biological membranes, gangliosides are found in conjunction with cholesterol, sphingomyelin and specific membrane proteins. Cholesterol is well known to modulate glycolipid conformation and enable receptor binding^42,75-79^. Here, we show that also the interaction between different gangliosides (*i*.*e*., NeuAc and NeuGc GM3, Figure 6D) can influence the conformation of glycolipids in the membrane – and make the gangliosides amenable to recognition by 14F7. This is valuable information, for two reasons: For one, it allows the detection of NeuGc GM3 even at low concentrations. This is the case in human cells (and although NeuGc GM3 content appears to increase in certain cancers, it is unclear if high concentrations can be reached). In contrast, the non-binding NeuAc GM3 is naturally expressed in all cells and may thus potentiate the recognition of low concentrations of NeuGc GM3 in cellular membranes, exposing the antigen for specific targeting. So far, we have only studied 14F7’s capacity to recognize NeuGc GM3 clusters in a membrane mimetic system, however, it is reasonable to assume that a similar effect occurs in a cellular environment. Although further investigation is certainly required, this would open up for completely new applications of 14F7, in the clinical setting as well as for biochemical analysis.

## Conclusions

We set out to characterize the exquisite specificity of 14F7 for NeuGc GM3, and its selectivity over NeuAc GM3, and have now solved the crystal structure of this promising anti-tumor antibody in complex with its target antigen by high-resolution X-ray crystallography. Complementary qualitative and quantitative liposome interaction studies with NeuGc GM3 additionally yielded unique insights into the formation and characteristics of glycolipid clusters and, potentially, membrane nanodomains.

Generally, antigen and immunogen are thought to have identical structures. Our data suggest that NeuGc GM3 concentration in the membrane influences its presentation and, in consequence, strongly affects recognition by 14F7. Therefore, although the immunogen is NeuGc GM3, the actual antigen recognized by 14F7 antibody is “high density NeuGc GM3”. Similar results were previously described for the M2590 mAb for NeuAc GM3^64^. Here we show additionally that different gangliosides can conformationally modulate each other, *i*.*e*., the presence of NeuAc GM3 helps convert NeuGc GM3 to its antigenic form. The switch between conformations appears to be abrupt, as caused by a phase transition, at sufficiently high concentration. However, binding requires that the membrane surface is relatively flat, as in GUVs and MLVs, and probably in biological membranes. We suspect that high positive curvature may disrupt the alignment of the saccharide head groups, abrogating binding of 14F7 (Figure 6). It may also dilute the concentration of negative charges, affecting electrostatic membrane-antibody interactions.

Our findings further suggest that under favorable conditions, such as the presence of interacting molecules like NeuAc GM3, NeuGc GM3 can be recognized even at relatively low concentrations in cellular membranes, despite requirement of a “high-density-form”. Our data thus inform new concepts for designing new immunotherapy strategies targeting glycolipids.

## Methods

### Synthesis of NeuGc trisaccharide

The NeuGc GM3 trisaccharide was synthesized through an IBr/AgOTf-promoted glycosylation of a benzylated lactose acceptor with a NeuGc thioglycoside donor, followed by global deprotection of the obtained trisaccharide as reported earlier^9^.

### Expression and purification of 14F7 derived scFv

The 14F7 scFv was produced by a variation of a protocol described by Bjerregaard-Andersen *et al*.^9^. Compared to the original 14F7 mAb, this construct contains an alternative light chain identified by Rojas *et al*.^8^. The linker was chosen on the basis of a vector system established for expression of single chain T-cell receptors (TCRs) and single-chain variable fragments (scFvs) in *Escherichia coli*^80,81^. Briefly, the scFv was expressed in *E. coli* by a pFKPEN vector-based system. The vector encodes a pelB leader sequence, thus promoting the translocation of the protein to the periplasm. Purification included limited lysis of the *E. coli* outer membrane to release the mature scFv, and subsequent purification by protein L affinity chromatography and size exclusion chromatography to reach a highly pure and homogenous preparation for crystallization and binding experiments.

### Crystallization of the 14F7 scFv in complex with NeuGc trisaccharide

Crystallization of the 14F7 scFv was performed as described earlier^9^. Crystals of good diffraction quality were obtained from the Morpheus screen (Hampton Research, US) after seeding with small crystals from initial hits. Remaining crystals from the D12 condition (12.5 % w/v PEG 1000, 12.5 % w/v PEG 3350, 12.5 % v/v MPD, 0.02 M 1,6-hexandiol, 0.02 M 1-butanol, 0.02 M(*RS*)-1,2-propanediol, 0.02 M 2-propanol, 0.02 M 1,4-butanediol, 0.02 M 1,3-propanediol, 0.1 M Bicine/Tris base pH 8.5), used for determination of the 14F7 scFv apo-structure^9^, were soaked by the addition of the synthesized NeuGc trisaccharide in powder form. The crystals were incubated for 1h before flash-cooling in liquid nitrogen and stored for diffraction experiments.

### Data collection and structure determination

Diffraction data extending to 2.3 Å were collected at the ID30A-3 beam line at the European Synchrotron Radiation Facility (ESRF), Grenoble, France. X-ray data were auto-processed at the ESRF by the EDNA pipeline^82^. The structure was phased by molecular replacement with the PHENIX crystallographic software package^83^, using the 14F7 scFv apo-structure (PDB ID: 6FFJ^9^) as search model, and refined in alternating cycles of manual model building and refinement with PHENIX^83^ and COOT^84^. Water molecules were built in at late stages of the refinement, initially using the automated finding by PHENIX. These sites were then inspected individually and assessed for removal in case of electron density sigma level >1.10 e/Å^3^ or bond distances >3.5 Å or <2.2 Å. Likewise, missing water molecules were added manually. The phased map revealed additional electron density in one of the four scFv molecules in the asymmetric unit, which was modelled as NeuGc GM3 trisaccharide. The trisaccharide ligand was built using eLBOW^85^ and modeled into the electron density of the binding pocket at final stages of structure building and adjusting occupancy by matching ligand *B*-factors to interacting protein residues. An OMIT difference density map was made by removing the trisaccharide ligand from the final model, followed by five refinement cycles using PHENIX^83^ The final model was deposited in the Protein Data Bank with accession code 6S2I.

### Modeling

The program VMD^86^ was used to for visualization and analysis, as well as for molecular modeling. The two amino acid substitutions in the heavy variable domain – S31R and W33Q – were made using the Mutator plugin implemented in VMD. Side chain conformations were modeled using the Molefacture plugin. This same tool was used to model the anticlinal conformation of the GM3 trisaccharide, keeping the sialic acid in its crystal position and modifying only the two torsion angles of its glycosidic linkage with galactose.

### Preparation of giant unilamellar vesicles (GUVs)

1,2-dioleoyl-*sn*-glycero-3-phosphocholine (DOPC) and cholesterol (both from Avanti, US) stocks were dissolved in chloroform/methanol (10:1), *N*-Acetyl (Matreya, US) and/or *N*-glycolyl GM3 (isolated from horse erythrocytes^87^ and kindly provided by CIM, Havana, Cuba) were dissolved in chloroform/methanol/water (2:1:0.1). GUVs were prepared by the polyvinyl alcohol (PVA; Mw 146,000-186,000, Sigma Aldrich, Germany) assisted method in the absence of divalent ions. Black, 96-well glass bottom plates (Greiner Bio-One) were coated with 2 % PVA and evaporated by heating at 70 °C for 20 min. In total 10 μl of desired lipid mixtures at 1 mg/mL in chloroform/methanol/water 1/2/0.8 (v/v) was spread on top of the PVA coated wells, followed by subsequent incubation under vacuum for at least 1 h to form a lipid film and allow removal of organic solvents. Sucrose containing swelling buffer (280 mM sucrose, 25 mM HEPES-NaOH, pH 7.4; 300 μl per well) was added and incubated for at least 20 min to induce vesicle formation.

### Confocal imaging

GUVs were deposited in a 1 % BSA precoated imaging chamber (384-Well Glass-Bottom Plates, Greiner Bio-One, Austria). Labeled 14F7 mAb (14F7hT) was added and the sample was left for 30 min incubation at 23 °C before confocal imaging. GUVs were imaged with a Zeiss LSM 780 confocal microscope. 488 nm and 543 nm lasers were used for excitation of green and red fluorophores, respectively. BP 530-550, BP 585-615 filters in multi-track mode were used to eliminate the cross talk.

### Preparation of liposomes

For vesicle preparation, lipids (DOPC and cholesterol from Avanti, US, NeuAc GM3 from Matreya, US, and NeuGc GM3 from CIM, Havana, Cuba) were mixed at the desired molar ratios and dried under nitrogen gas stream, followed by incubation under vacuum for 4 h to remove organic solvents. The dried lipid film was re-hydrated in HEPES-buffered saline (HBS) (10 mM HEPES -NaOH, 150 mM NaCl, pH 7.4) to a final concentration of 1 mg/mL, for 15 min at 600 rpm. Multilamellar vesicles (MLV) were either used directly or further processed to yield unilamellar vesicles (LUVs). LUVs where produced by subjecting the liposomes to 10 cycles of freezing in liquid nitrogen and subsequent thawing in a heating block at 30 °C. The vesicle solution was extruded 21 times through a 100 nm diameter polycarbonate membrane (Whatman® Nuclepore, Fisher Scientific, US) using an extrusion kit (Avanti, US). The size of the liposomes was determined by dynamic light scattering using a Zetasizer Nano ZS (Malvern Instruments, UK).

### Lipid extraction and validation by thin-layer chromatography (TLC)

Lipid composition of liposomes was assayed by thin layer chromatography as described previously^88^. Briefly, lipids were extracted using two step extraction protocol (chloroform:methanol 10:1 followed by 2:1)^88^. After each step, the lipid containing organic phase was pooled and dried under a nitrogen stream. The lipids were resuspended in a small volume of chloroform/methanol (2:1) and applied to a to HPTLC plate (Silica Gel 60, Merck, Germany) together with DOPC/cholesterol and NeuGc/NeuAc GM3 as standards. The plate was placed under vacuum for 30 min. For development, chloroform/methanol/ 0.2 % calcium chloride (60/35/6) was used. To visualize glycolipids, the plate was sprayed with orcinol (Sigma Aldrich, Germany) and heated (200 °C until desired signal was observed).

### Flotation assay

50 µl of liposomes was mixed with 14F7 mAb (14F7hT) or scFv (20 µg/mL final concentration) and incubated for 30 min on ice. Thereafter, iodixanol (Optiprep; Sigma Aldrich, Germany) was added to a final concentration of 30 % and a step gradient was built on top (10 %, 2.5 %, and 0 % iodixanol in HBS). Protein-liposome complexes were separated from unbound protein by centrifugation (2 h at 45.000 rpm, 4 °C, MLS50 rotor -Beckman Coulter) in the density gradient. After flotation, 11 fractions were collected from top of the tube. Samples were precipitated with 10 % TCA and pelleted by centrifugation (20 min, 20,000 x *g*, 4 °C). The supernatant was discarded, and the remaining pellet was neutralized with 1.5 M Tris -HCl, pH 8.8. SDS loading buffer (250 mM Tris-HCl pH 6.8, 12.5 mM EDTA, 10 % SDS, 25 % glycerol, 200 mM DTT) was added to all samples before loading on a 4-12 % Bis-Tris gel for SDS-electrophoresis.

### Electrochemiluminescence immunoassay (EIA)

Liposomes were passively adsorbed on the electrode surface (1 h, 23 °C), and the residual sites on the surface were blocked with 0.2 % porcine gelatin (1 h, 23 °C). The surface was then washed three times with HBS and porcine gelatin solutions containing the desired concentrations of 14F7 mAb or scFv (were added to each well). Binding was carried out for 2 h at 23 °C. Wells were then washed and a solution of goat anti-human-SulfoTAG (Mesoscale Discovery, US) or protein L-SulfoTAG was added (1 µg/mL, 23 °C, 1 h). The wells were washed and reading buffer was added (MSD surfactant-free reading buffer). The background was determined from binding of secondary antibodies or protein L to liposomes. Data were acquired on a SECTOR Imager 6000. The recorded data were analyzed using GraphPad Prism 6.0 software using one site-specific binding algorithm.

### Labeling of humanized 14F7 mAb (14F7hT) and protein L

14F7hT was labeled using the DyLight 488-NHS ester (Thermo Fisher, US) for detection during confocal microscopy. Protein L was labeled with SulfoTAG-NHS ester (Mesoscale Discovery, US) at a 1:20 molar ratio for scFv detection during EIA. The mixture was incubated for 1 h at 4 °C. The reaction was stopped by the addition of at least 5-fold molar excess of ethanolamine over the NHS-reagent, followed by the removal of excess label-molecules by a gravity spin column (GE Healthcare, Germany).

### Thermal stability measurements using nanoDSF

For nanoDSF measurements, scFv was diluted with HBS to reach a final concentration of 0.5 mg/mL, and subsequently filled into nanoDSF standard treated capillaries. Thermal unfolding and aggregation was monitored in a temperature ramp with 1 °C/min from 20 °C to 95 °C with a resolution of ∼ 20 data points/min. Analysis of unfolding and aggregation was performed using the PR.Control Software.

### Surface Plasmon Resonance

SPR experiments were performed using a BIAcore X100 instrument on an HPA sensor chip^89^ (GE Healthcare, Germany). An HPA chip was cleaned with 40 mM octyl-glucoside (10 µL/min, 5 min). Next, LUVs consisting of DOPC, cholesterol (both from Avanti, US), NeuGc GM3 (CIM, Havana, Cuba) and/or NeuAc GM3 (Matreya, US) at the desired molar ratios, were injected across the sensor chip at a low flow rate. The LUVs (400 µM) were allowed to collapse and form a monolayer on the HPA chip (2 µL/min, 15 min). Then, the chip was washed twice with 10 M NaOH (10 µL/min, 30 s), and once with HBS (10 mM HEPES, 150 mM NaCl, pH 7.4) (80 ul/min, 60 s) to remove any unbound liposomes. The end-response was between 900 and 1300 RU. 14F7 mAb (500 nM) in HBS was injected at 30 µL/min for 3 min. The dissociation phase was measured for 5 min. At the end of the binding assay, the surface of the sensor chip was regenerated with two injections of 3:2 10 M NaOH/isopropanol (10 μL/min, 30 s). The binding response of the 14F7 mAb was obtained after subtracting the binding signal from the reference flow cell containing vesicles without gangliosides. The experiment was repeated twice with similar results.

### Western blot

Proteins were transferred to a 0.45 *μ*m membranes (Millipore) for immunoblotting. The membrane was incubated for 1 h on a tilting tray (10 rpm, 23 °C) in TM-PBS buffer (0.1 % Tween 20, 5 % nonfat dry milk powder in 1x PBS), before a 20 sec wash in fresh TM-PBS. Secondary antibody (Goat anti-human IgG, 1.25 *μ*g/mL, SulfoTAG) diluted in TM-PBS was added, and the membrane was incubated for 1 h (10 rpm, 23 °C). The membrane was washed thrice in 5 min intervals, twice in T-PBS (0.1 % Tween 20 in 1x PBS) and once in 1x PBS. Detection was performed with a CCD imager (Imager 600, GE Healthcare) using SuperSignal West Pico PLUS (Thermo Fisher, US) as substrate.

## Acknowledgments

We thank the staff of ESRF (Guillaume Gotthard) for assistance and support in using beamline ID30A-3, and the Center of Molecular Immunology (CIM), Havana for providing us with 14F7 mAb and the NeuGc GM3 ganglioside. Work at UiO was funded by the University of Oslo (including the postdoc position of KBA and the PhD position of HJ). Work in Dresden was supported by the German Federal Ministry of Education and Research (BMBF) grant to the German Center for Diabetes Research (DZD e.V.; Ü.C.) and the Deutsche Forschungsgemeinschaft (DFG; Project Number 251981924 – TRR 83; Ü.C.). Work at UdeM was supported with funds from the Colombian Government (NanoBioCáncer program, grant FP44842-211-2018), and work at UCD was funded by Science Foundation Ireland, grants No. 08/SRC/B1393 and13/IA/1959. Work at NIC was supported by the Slovenian Research Agency (program grant P1-0391).

## Author contributions

H.J., M.G., Ü.C. and U.K. conceived the study. F.A. synthesized the trisaccharide, supervised by S.O.. H.J. expressed and purified the constructs, supervised by U.K.. K.B.-A. was in charge of the crystallography, with U.K. validating the crystal structure. E.M. performed the modeling studies. Liposome experiments were performed by H.J., D.G. and M.G., who also served as supervisor for this part of the work. SPR experiments were performed by H.J. and A.S., supervised by G.A.. K.B.-A. and H.J. wrote the first draft of the manuscript, which was revised in tight collaboration with E.M., M.G. and U.K., and approved by all authors.

## Supplementary Information

**Figure S1.**
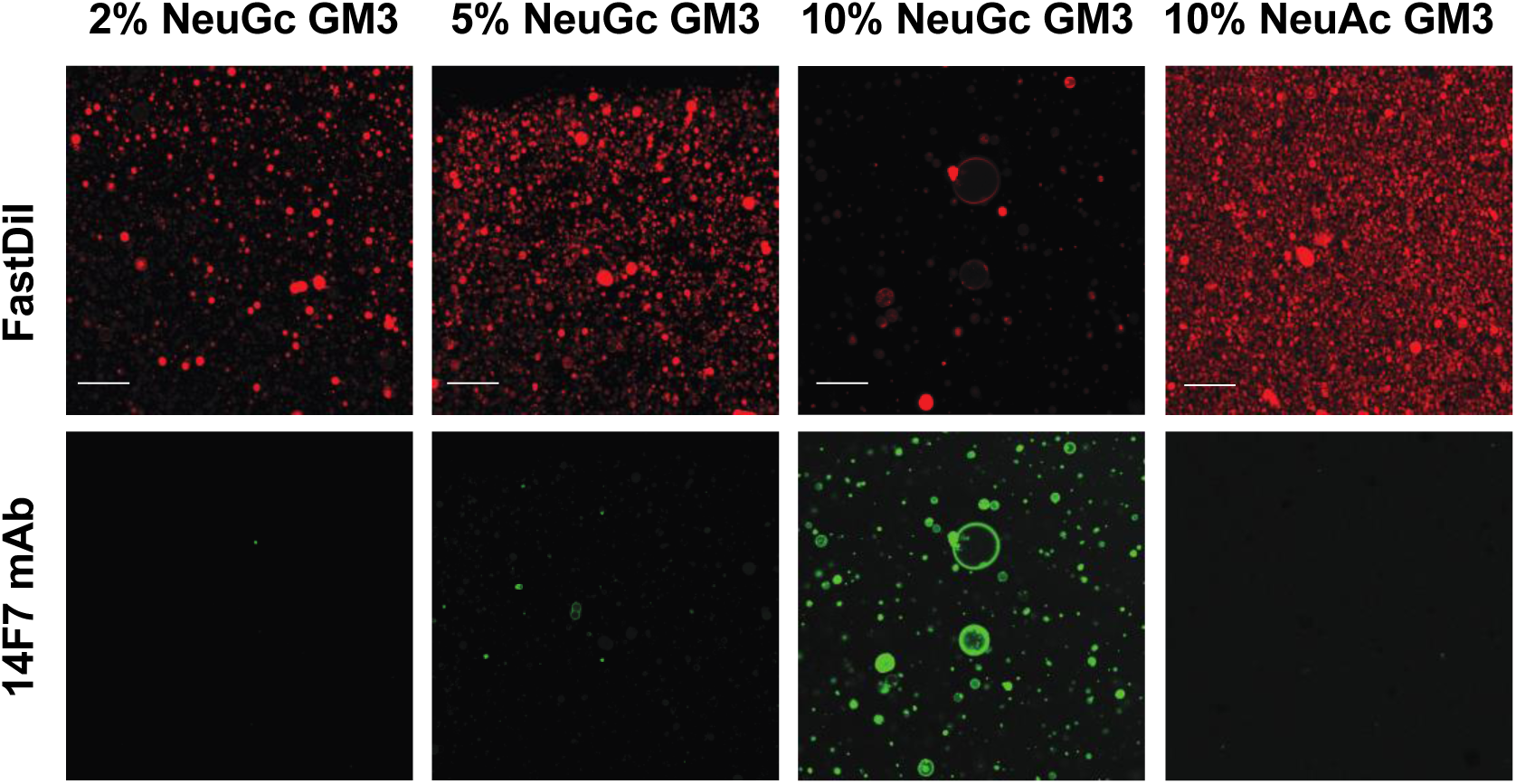
Binding of 14F7 to GUVs composed of DOPC/Chol/GM3. 14F7 was chemically labeled with Dylight-488 (green) and added to GUVs containing FastDiI as a membrane marker (red). While there was negligible interaction between 14F7 mAb and GUVs with 2 % and 5 % NeuGc GM3, 14F7 bound strongly to GUVs containing 10 % NeuGc GM3. No 14F7 mAb binding was observed for GUVs with 10 % NeuAc GM3. The scale bar corresponds to 20 μm.

**Figure S2.**
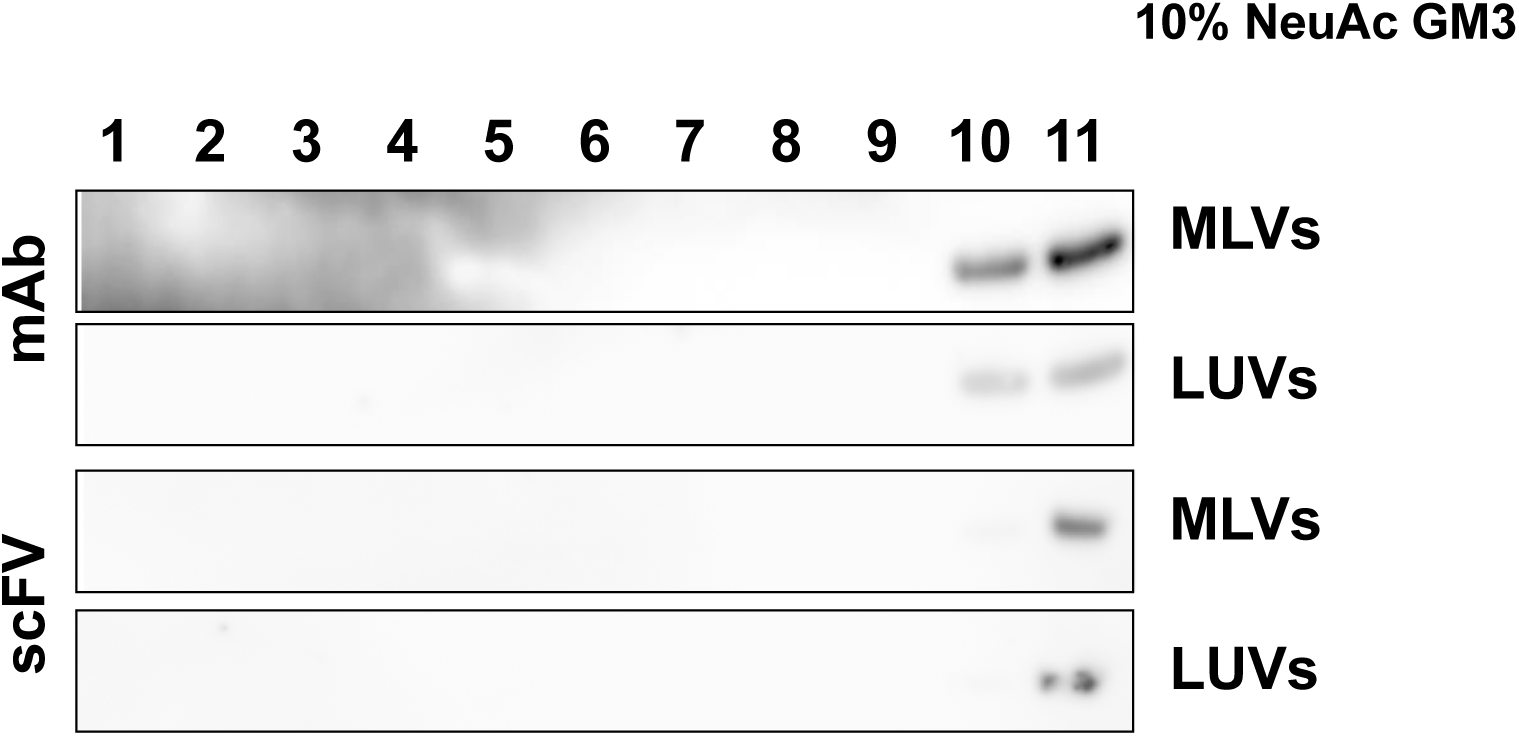
Binding of 14F7 to LUVs and MLVs composed of 10 % NeuAc GM3/DOPC/ cholesterol in a flotation assay. After centrifugation, 11 fractions were collected from the top. Proteins bound to vesicles accumulated in fractions 2-4. For all flotation assays with 10 % NeuAc GM3, the protein was located in the bottom fractions (10-11), where unbound proteins accumulate.

**Figure S3.**
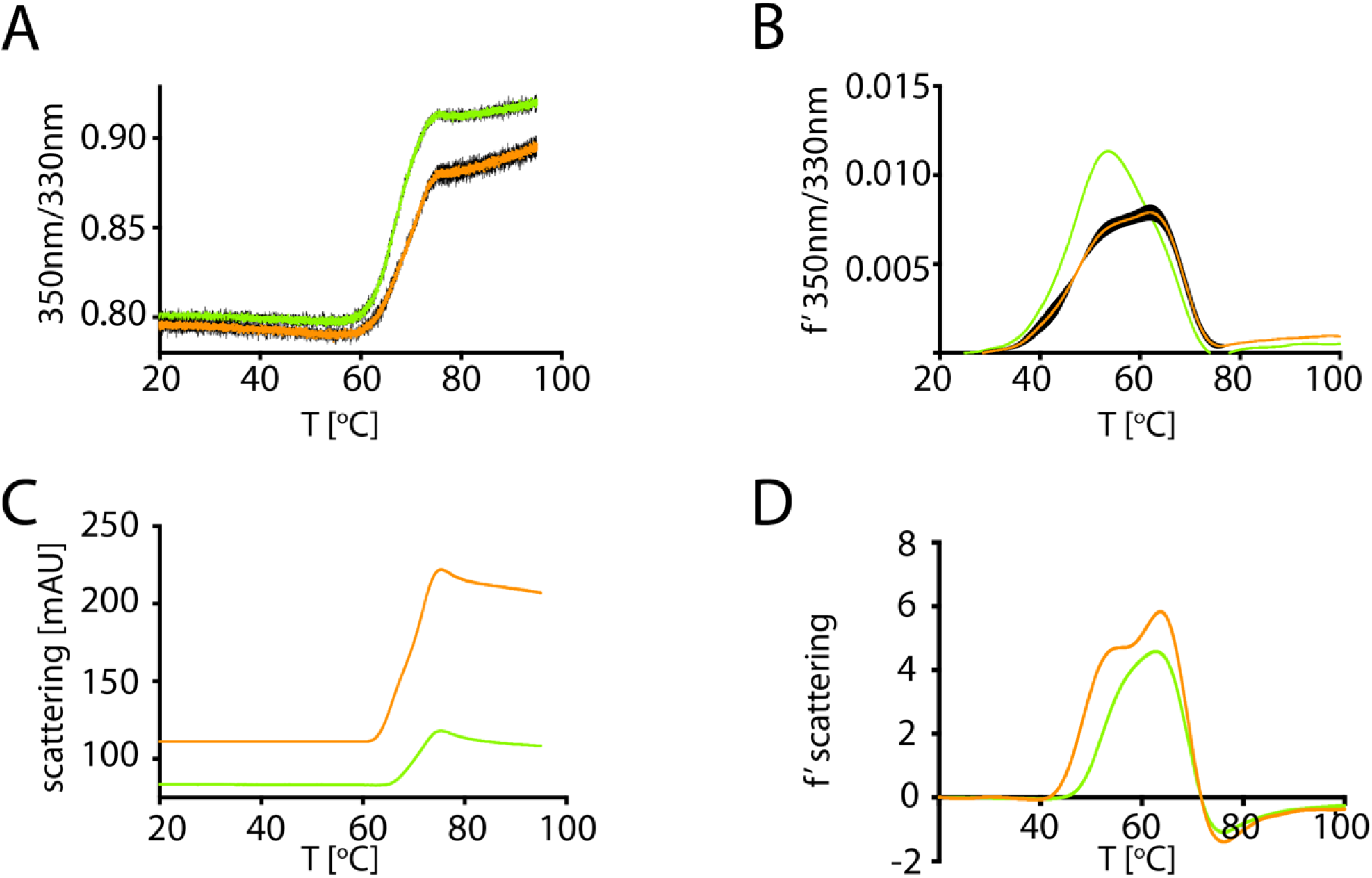
nanoDSF measurements of native (green) and Dylight488-labeled (orange) 14F7 scFv. **A** Thermal unfolding curves. **B** First derivative of unfolding curves. **C** Aggregation propensity curves. **D** First derivative of aggregation propensity curves.

## Notes

### Competing Interest Statement

The authors have declared no competing interest.

